# Methionine stress induces a ferroptotic gene signature in methionine dependent cancer cells

**DOI:** 10.1101/2020.08.18.254490

**Authors:** Katherine Wallis, Jordan T. Bird, Allen Gies, Sam G. Mackintosh, Alan J. Tackett, Stephanie Byrum, Isabelle R. Miousse

**Affiliations:** Department of Biochemistry and Molecular Biology, University of Arkansas for Medical Sciences, Little Rock, AR; Bioinformatics Core, University of Arkansas for Medical Sciences, Little Rock, AR

**Keywords:** Methionine, ferroptosis, proteomics, melanoma, OXPHOS

## Abstract

Dietary methionine restriction is associated with a reduction in tumor growth in preclinical studies and an increase in lifespan in animal models. The mechanism by which methionine restriction inhibits tumor growth while sparing normal cells is incompletely understood, except for the observation that normal cells can utilize methionine or homocysteine interchangeably (methionine independence) while most cancer cells are strictly dependent on methionine availability. Here, we compared a typical methionine dependent and a rare methionine independent melanoma cell line. We found that replacing methionine with homocysteine generally induced hypomethylation in gene promoters. We isolated nuclear proteins and submitted it for tandem mass tag (TMT) proteomics. This analysis revealed that several proteins involved in the mitochondrial integrated stress response (ISR) were upregulated in response to the replacement of methionine to homocysteine in both cell lines, but to a much greater degree in the methionine dependent cell line. Consistent with the ISR signature, a proteomic analysis of a subcellular fraction enriched for mitochondrial content revealed a strong enrichment for proteins involved in oxidative phosphorylation. Analysis of cellular bioenergetics confirmed that homocysteine induces a decrease in ATP production from oxidative phosphorylation and glycolysis, but to a similar extent in methionine dependent and methionine independent cells. The mitochondrial integrated stress response shared a signature with ferroptosis. Methionine dependent cells displayed a strong ferroptotic signature, which was decreased by half in methionine independent cells. Consistent with ferroptosis, lipid peroxidation was significantly increased in methionine independent cells grown in homocysteine, and viability could be rescued partially but significantly with the inhibitor ferrostatin. Therefore, we propose that methionine stress induces ferroptotic cell death in methionine dependent cancer cells.

## INTRODUCTION

Fundamental differences in metabolism between normal and cancer cells represent optimal targets for cancer therapy, minimizing the risks for adverse effects. One such fundamental difference was identified over 40 years ago in the way normal cells and cancer cells respond to methionine depletion, and more particularly in their response to methionine depletion when the intermediate homocysteine is provided (1, 2). Most normal cells experience a small and slow decrease in proliferation under methionine depletion. Addition of homocysteine to the culture media completely rescues proliferation. Cancer cells, on the other hand, die within 48-72h after the initiation of methionine depletion, and rescue by homocysteine is modest (1). Exceptions exist to this rule. Activated T cells are not rescued by homocysteine (3). Conversely, rare cancer cell lines, such as the human melanoma MeWo, can be rescued nearly completely with homocysteine (3). *In vivo*, enzymatic and dietary methionine depletion reduces tumor growth and synergizes with many cancer therapies (5–9). At the same time, dietary methionine restriction is associated with improvements in glucose and lipid regulation, and in a general increase in lifespan (10). However, the nature of the difference in response to methionine depletion between normal cells and cancer cells remains unknown.

Methionine is an essential amino acid fulfilling several crucial metabolic roles. In addition to protein synthesis, methionine is the precursor for the universal methyl donor S-adenosylmethionine. S-adenosyl methionine donates methyl groups to several acceptor molecules such as DNA, RNA, proteins (including, but not restricted to, histones), and phospholipids. The resulting demethylated molecule, S-adenosylhomocysteine, yields homocysteine which can then be either enter the remethylation pathway to methionine or the transulfuration pathway to the antioxidant glutathione. Alternatively, S-adenylsylmethionine can be decarboxylated and enter the methionine salvage pathway, resulting in the production of polyamines.

While methionine synthase activity is similar between methionine dependent and independent cell lines, the abundance of methyl donors is different between the phenotype (11). Treatment with an inhibitor of DNA methyltransferase was also found to cause a reversion (11). Comparison of the methionine independent MeWo cell line with its more aggressive, methionine dependent variant MeWo-LC1 identified that hypermethylation of the promoter region of the vitamin B12 chaperone MMACHC explained the divergence in methionine phenotype (4). However, this finding was limited to this single cell line. The use of methyl groups was also reported to be different between methionine dependent and independent cells. The methyl group from methionine is used primarily for the synthesis of S-adenosylmethionine and methylated proteins in methionine independent cells (12). Methionine dependent cells, on the other hand, primarily use this methyl group for the synthesis of creatine and phosphatidylcholine.

In this study, we sought to understand the many ramifications of the replacement of methionine with homocysteine in cell culture media. We compared the methylome and proteome of two human melanoma cell lines. One, A101D, is a prototypical cancer cell line displaying methionine dependence. The second, MeWo, has long been described as methionine independent, i.e. rescued by homocysteine. We identified a general decreased in promoter DNA methylation, with changes in the nuclear proteome suggesting the activation of the mitochondrial integrated stress response. Changes in the mitochondrial proteome pointed to alterations in oxidative phosphorylation. Cellular bioenergetics did indicate a decrease in cellular respiration, but this response was not different between methionine dependent and independent cells. However, a ferroptosis signature pointed to a divergence in lipid peroxidation between methionine dependent and independent cells.

## MATERIALS AND METHODS

### Cell culture

A101D and MeWo cells were obtained from ATCC. Cells were cultured in high glucose, no glutamine, no methionine, no cystine DMEM (ThermoFisher Scientific, Waltham MA) supplemented with 5% dialyzed serum (BioTechne, Minneapolis, MN), 100 IU penicillin and streptomycin (ThermoFisher), 4 mM L-glutamine (ThermoFisher) and 1 mM sodium pyruvate (ThermoFisher). L-cystine (Millipore-Sigma, Burlington, MA) was resuspended in PBS with NaOH added until complete solubilization, and added to the cell media at a final concentration of 150 μM. For control medium, L-methionine (Millipore-Sigma) was resuspended in PBS and added to the cell media at a final concentration of 200 μM. For methionine-free, homocysteine containing media, L-homocysteine (Chem-impex, Wood Dale, IL) was resuspended in PBS with NaOH added until complete solubilization, and added to the cell media at a final concentration of 200 μM.

### Cell cycle

A101D and MeWo cells were cultured in either methionine or homocysteine media for 24 hours or 48 hours. Cells were fixed in ethanol, washed, and treated with ribonuclease. Finally, cells were stained with DAPI and analysed for cell cycle by fluorescence activated sorting on a BD LSRFortessa instrument (Beckman Coulter, Brea, CA).

### DNA methylation

A101D and MeWo cells were grown in either control media or homocysteine media for 24 hours. DNA and RNA was extracted using the AllPrep kit following the manufacturer’s instructions (Qiagen, Germantown, MD). After Qubit fluometric quantification (ThermoFisher), methylation was analyzed using the Infinium MethylationEPIC array (Illumina, San Diego, CA) at the Genomics Core of the University of Arkansas for Medical Sciences. Data analysis was performed at the Bioinformatics Core. Methylation was expressed as the absolute difference between methionine and homocysteine.

DNA methylation data was preprocessed and normalized using Bioconductor packages *minfi* and *watermelon* (13,14). Briefly, samples were corrected for background fluorescence and dye biases and poor CpGs were identified and removed. Samples where more than 10% of probes had detection p-values >1×10^−5^ or had irregular intensity distributions were excluded. The resulting data was then normalized using the Beta-Mixture Quantile Normalization method (15).

Differential DNA methylation was analyzed using the R statistical programming language (version 3.6) and the Bioconductor package *Limma* (16). Each CpG’s p-value was adjusted for multiplicity using the Benjamini and Hochberg method (17) to control the False Discovery Rate (FDR) and CpGs with FDR <0.05 were accepted as statistically significant. Probes with a median detection p-value >0.05 were excluded from analysis. Pathway analysis was performed using the Bioconductor package *missMethyl* (18).

### Subcellular protein fractions

A101D and MeWo cells were grown in either control media or homocysteine media for 24 hours. Nuclear proteins were extracted using the EpiQuik Nuclear Extraction Kit following the manufacturer’s instructions (Epigentek, Farmingdale, NY). Nuclear extracts were frozen at − 80°C until use. A crude mitochondrial fraction was obtained as described by Graham et al. (19). Briefly, cells were homogenized in general purpose buffer (0.25 M sucrose, 1 mM EDTA, 10 mM Hepes-NaOH, pH 7.4) and centrifuged at 1,000g for 10 min. The crude nuclear pellet, inferior in quality to the one described above, was discarded and the supernatant was transferred to a fresh tube and further centrifuged at 16,000g for 10 min. Crude mitochondrial pellets were stored at −80°C until use.

### Proteomics

Purified proteins were reduced, alkylated, and digested using filter-aided sample preparation (20). Tryptic peptides were labeled using a tandem mass tag 10-plex isobaric label reagent set (ThermoFisher) following the manufacturer’s instructions. Labeled peptides will be separated into 36 fractions, and then consolidated into 12 super-fractions. Each super-fraction were further separated by reverse phase. Peptides were eluted and ionized by electrospray followed by mass spectrometric analysis on an Orbitrap Fusion Lumos mass spectrometer (ThermoFisher). MS data were acquired using the FTMS analyzer and MS/MS data using the ion trap analyzer. Up to 10 MS/MS precursors were selected for HCD activation, followed by acquisition of MS3 reporter ion data. Proteins were identified and reporter ions quantified by searching the *Homo sapiens* (February 2020) UniprotKB database using MaxQuant (version 1.6.10.43, Max Planck Institute) with a parent ion tolerance of 3 ppm, a fragment ion tolerance of 0.5 Da, and a reporter ion tolerance of 0.001 Da. Protein identifications were accepted if they can be established <1.0% false discovery and contained at least 2 identified peptides. Protein probabilities were assigned by the Protein Prophet algorithm (21). Protein TMT MS3 reporter ion intensity values were assessed for quality using our in-house ProteiNorm app, a user-friendly tool for a systematic evaluation of normalization methods, for comparison of different differential abundance methods. Popular normalization methods were evaluated including log2 normalization (Log2), median normalization (Median), mean normalization (Mean), variance stabilizing normalization (VSN) (22), quantile normalization (Quantile) (23), cyclic loess normalization (Cyclic Loess) (16), global robust linear regression normalization (RLR) (24), and global intensity normalization (Global Intensity) (24). The individual performance of each method was evaluated by comparing of the following metrices: total intensity, Pooled intragroup Coefficient of Variation (PCV), Pooled intragroup Median Absolute Deviation (PMAD), Pooled intragroup estimate of variance (PEV), intragroup correlation, sample correlation heatmap (Pearson), and log2-ratio distributions. The data was normalized using log2 Cyclic Loess for the nuclear samples and VSN was used for the mitochondrial samples since these methods had the lowest variance and the highest sample correlations for the data sets. The normalized data was used to perform statistical analysis using Linear Models for Microarray Data (limma) with empirical Bayes (eBayes) smoothing to the standard errors (16). Proteins with an FDR adjusted p-value <0.05 and a fold change >2 were considered to be significant. Significant proteins were utilized to identify important protein networks and pathways using the Ensemble of Gene Set Enrichment Analyses Bioconductor package and Qiagen’s Ingenuity Pathway Analysis (25).

### Cellular bioenergetics

To measure oxygen production and acidification resulting from cellular metabolism, we plated A101D and MeWo cells at 30,000 live cells per well in a 96-well plate (Agilent, Santa Clara, CA). Cells were plated in either control or homocysteine media and incubated for 24 hours. The next day, the cell media was replaced with the assay media and ran using the Seahorse XF Cell Mito Stress Test and ATP Rate Assay on a Seahorse XFe96/XF96 analyzer (Agilent). Hoechst 33342 staining (ThermoFisher) was used to confirm that differences were not due to a variation in cell number.

### RNA-Seq

RNA was extracted as described above. After Qubit fluometric quantification (ThermoFisher), the NGS library was prepared with the Truseq Stranded Total RNA (Illumina, San Diego, CA) and ran on a HiSeq300 instrument at the Genomics Core of the University of Arkansas for Medical Sciences. Data analysis was performed at the Bioinformatics Core. Briefly, the RNA reads were checked for quality of sequencing using FastQC. RNA reads were trimmed using Trimmomatic using the following parameters: LEADING:3 TRAILING:3 SLIDINGWINDOW:4:28 MINLEN:36. Reads that pass quality control were aligned to the reference genome (hg38.p19 Acc:GCA_000001405.28) with Ensembl GRch38 release 99 annotations using the STAR sequence aligner. The RNA counts were extracted using featureCounts (26) and filtered (counts per million < 10 or < 20 across all samples) normalized using trimmed mean of M-values (TMM) using functions with EdgeR (27,28). Read TMM were then prepared to fit linear models using voomWithQualityWeight (29) and differential expression analysis was performed using limma (16). Genes with an FDR p-value < 0.05 and a fold change > 2 were considered significant.

### Lipid Peroxidation

A101D and MeWo cells were plated at a cell density of 10,000 cells per well in a 96-well plate and let to attach overnight. The next day, media was changed for control media, control media supplemented with homocysteine, or methionine-free, homocysteine-supplemented media for 24 hours (N=8). Cell were washed in PBS then incubated with 10 μg/mL BODIPY dye (ThermoFisher) in PBS for 30 min at 37°C. Fluorescence intensity was measured after 3 PBS washes as a ratio of emission at 591nm to 581nm.

### Viability and rescue

To viability rescue with different compounds, we plated A101D and MeWo in a 96 well plate at 10,000 cells per well. Cells attached for 4-5 hours and media was replaced with methionine-containing or homocysteine-containing media. N-acetyl-L-cysteine, choline chloride, ferrostatin, and Z-VAD-FMK were purchased from Millipore-Sigma. We measured viability after 72h by fluorescence with resazurin at a final concentration of 10 μg/mL (ThermoFisher).

## RESULTS

### Methionine replacement with homocysteine maintains viability in MeWo but not A101D cells

To confirm the methionine dependence status of human melanoma cells MeWo and A101D, we performed a cell cycle analysis after 24 and 48 hours. After 24 hours, there was an increase in cells in the G1 phase and a decrease in cells in the G2/M phase in A101D, while the reverse was true in MeWo (Fig 1, A and B). After 48 hours, there was a large increase in SubG1 cells that was specific to A101D, while the increase in G2/M cells was still evident in MeWo (Fig 1, panels C and D). The presence of cleaved DNA in A101D confirms that cell death occurred under homocysteine conditions in these cells (methionine dependence), but not in MeWo (methionine independence).

**Figure 1.**
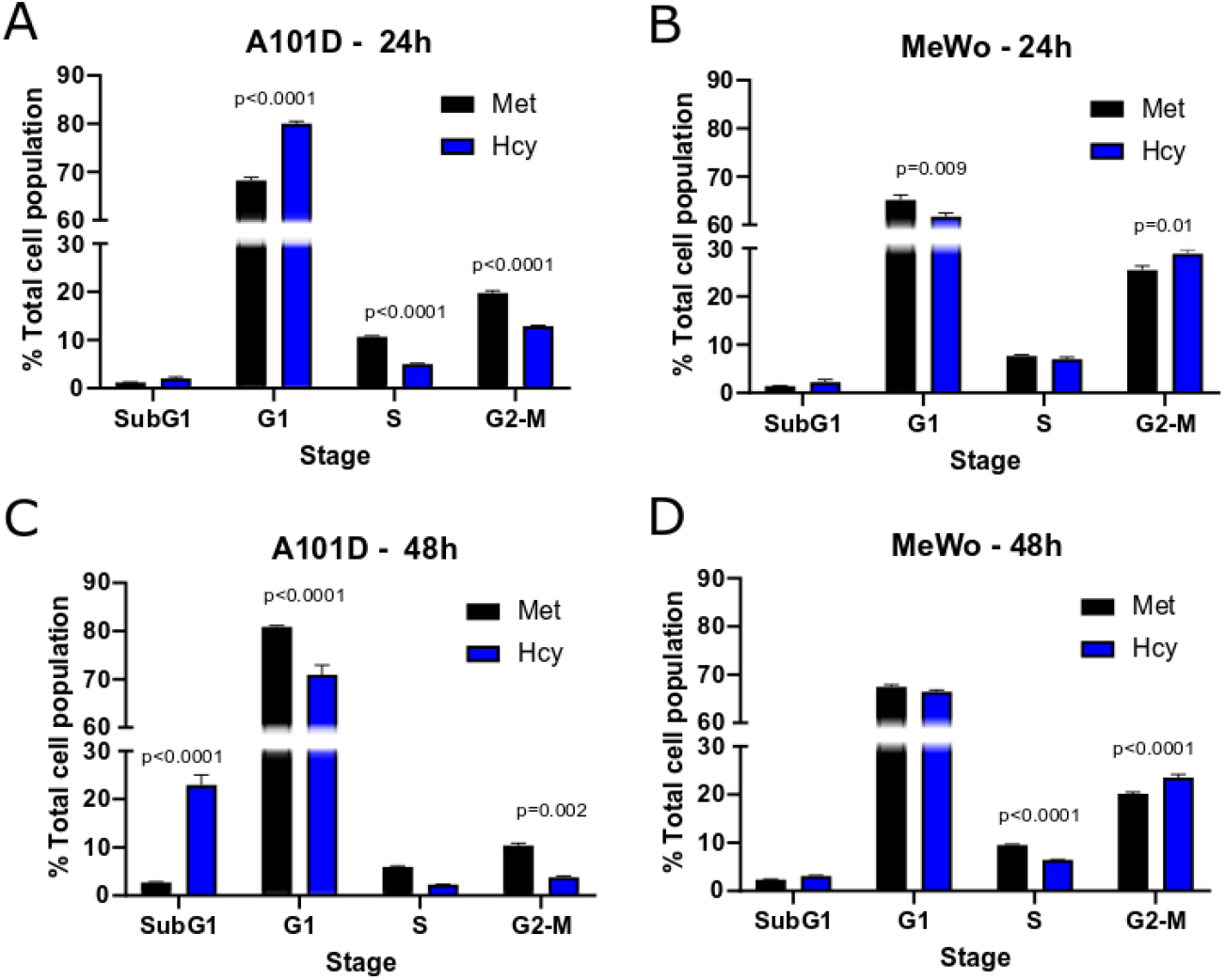
Cell cycle analysis in cells fed either methionine or homocysteine. A101D (Panels A and C) and MeWo (B and D) human melanoma cells were cultured in media containing methionine or methionine-free media supplemented with homocysteine for 24h (A and B) or 48h (C and D). Met: Methionine, Hcy: Methionine-free, homocysteine supplemented medium. Standard error of mean shown. P value calculated for T-test between Met and Hcy.

### Methionine replacement with homocysteine induces hypomethylation in both MeWo and A101D cells

Methionine is an important methyl donor in cells and we therefore investigated the DNA methylation patterns in methionine dependent and independent human melanoma cells cultured with either methionine or homocysteine. We chose a time point of 24 hours to measure cells prior to cell death. Beta values were significantly decreased in homocysteine versus methionine for both cell lines, indicating a net decrease in methylation levels (Fig. 2A). Interestingly, the beta value was decreased to a greater extent in MeWo than in A101D (Fig. 2C), despite the cell cycle phenotype being significantly milder. Accordingly, a total of 252 probes were significantly differentially methylated between methionine and homocysteine in the methionine dependent A101D, compared to 1145 in MeWo. All significant sites in both cell lines were hypomethylated in homocysteine compared to methionine. IPA analysis suggested that Sonic Hedgehog signaling was the main canonical pathway dysregulated in A101D and calcium signaling in MeWo (Fig 2B and D).

**Figure 2.**
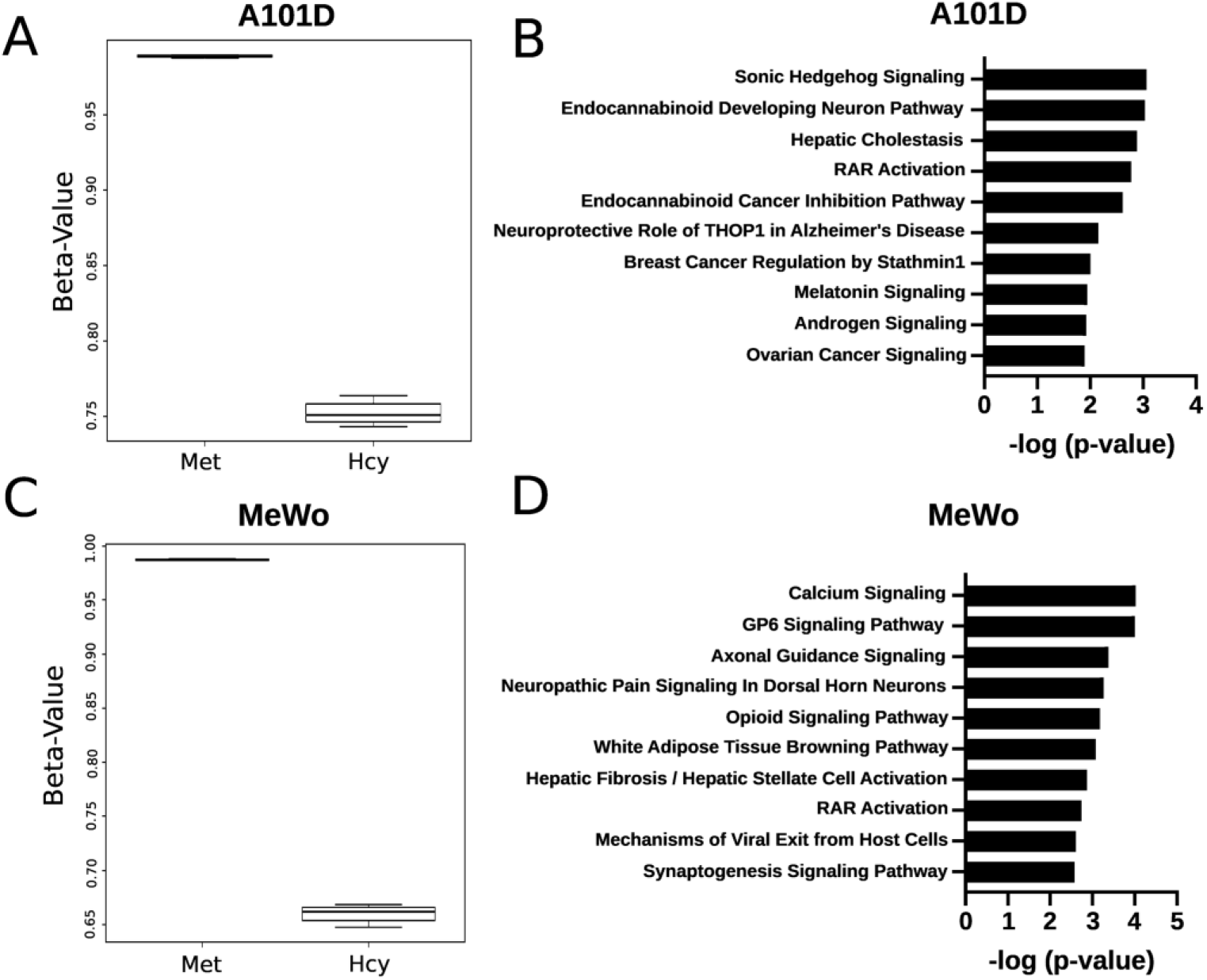
Whole genome methylation analysis in cells fed either methionine or homocysteine. Beta values representing an estimation of methylation levels for A101D (A) and MeWo (B) human melanoma cells cultured in media containing methionine or methionine-free media supplemented with homocysteine for 24h. Top 10 significantly enriched pathways for genes associated with differentially methylated sites in A101D (C) and MeWo (D) (IPA analysis). Met: Methionine, Hcy: Methionine-free, homocysteine supplemented medium.

### Methionine replacement with homocysteine is associated with a mitochondrial integrated stress response signature in the nuclear proteome

In order to investigate further the signaling events suggested by the methylation array, we analyzed the nuclear proteome of cell growing in either methionine or homocysteine. This approach allowed us to be sensitive to the translocation of proteins to the nucleus, even when transcription or total protein levels are constant. IPA pathway analysis revealed cholesterol biosynthesis in both cell lines, and telomerase and DNA damage pathway that were specific to A101D (Supplementary Fig 1). Upon closer analysis, the most enriched protein in homocysteine compared to methionine in A101D was ATF4 (12 fold, Table 1). ATF4 is a major transcription factor involved in the mitochondrial integrated stress response (mISR). Under stress, translation is paused except for the preferential translation of ATF4, allowing for the transcription of ATF4 target genes. We identified several other enriched protein related to the mISR such as DDIT3 (CHOP; 5 fold), CEBPB (3.6 fold), ATF3 (3 fold), and SESN2 (2.4 fold). An enrichment in ATF4 targets was also detected in MeWo with significant increases in DDIT3 (4.2 fold), SESN2 (2.9 fold), CEBPB (2.2 fold), as well as in the protein NUPR1 (3.5 fold). These results suggest that the replacement of methionine by homocysteine induces the mISR in both cell lines.

**Table 1.** ATF4 and ATF4-related proteins.

|  | **A101D** | **MeWo** |
| --- | --- | --- |
| **ATF4** | 12.13 | undet |
| **DDIT3** | 5.28 | 4.29 |
| **CEBPB** | 3.03 | 2.14 |
| **ATF3** | 3.03 | undet |
| **SESN2** | 2.46 | 2.83 |
| **ASNS** | 1.51 | 1.34 |
| **PPP1R15A** | undet | 2.66 |
| **NUPR1** | undet | 3.43 |

### Proteins involved in oxidative phosphorylation are enriched in methionine dependent cells

The mISR signature in the nuclear proteome suggested important mitochondrial effects in response to homocysteine versus methionine. This led us to analyze a crude mitochondrial protein extract. This extract significantly enriches for mitochondrial proteins along with ER and Golgi, following a simple procedure and leading to high yields. The top dysregulated pathway in both A101D and MeWo was “EIF2 Signaling”, confirming our nuclear results with the ATF4 pathway (Fig 3, A and B). Interestingly, the analysis predicted that the regulation was in opposite direction in the two cell lines; downregulated in A101D and upregulated in MeWo. The pathway “Oxidative Phosphorylation” was upregulated in A101D, but only modestly in MeWo. An EGSEA analysis showed that “Oxidative Phosphorylation” was the most upregulated hallmark in A101D and did not appear in MeWo (Figure 3 panels C and D).

**Figure 3.**
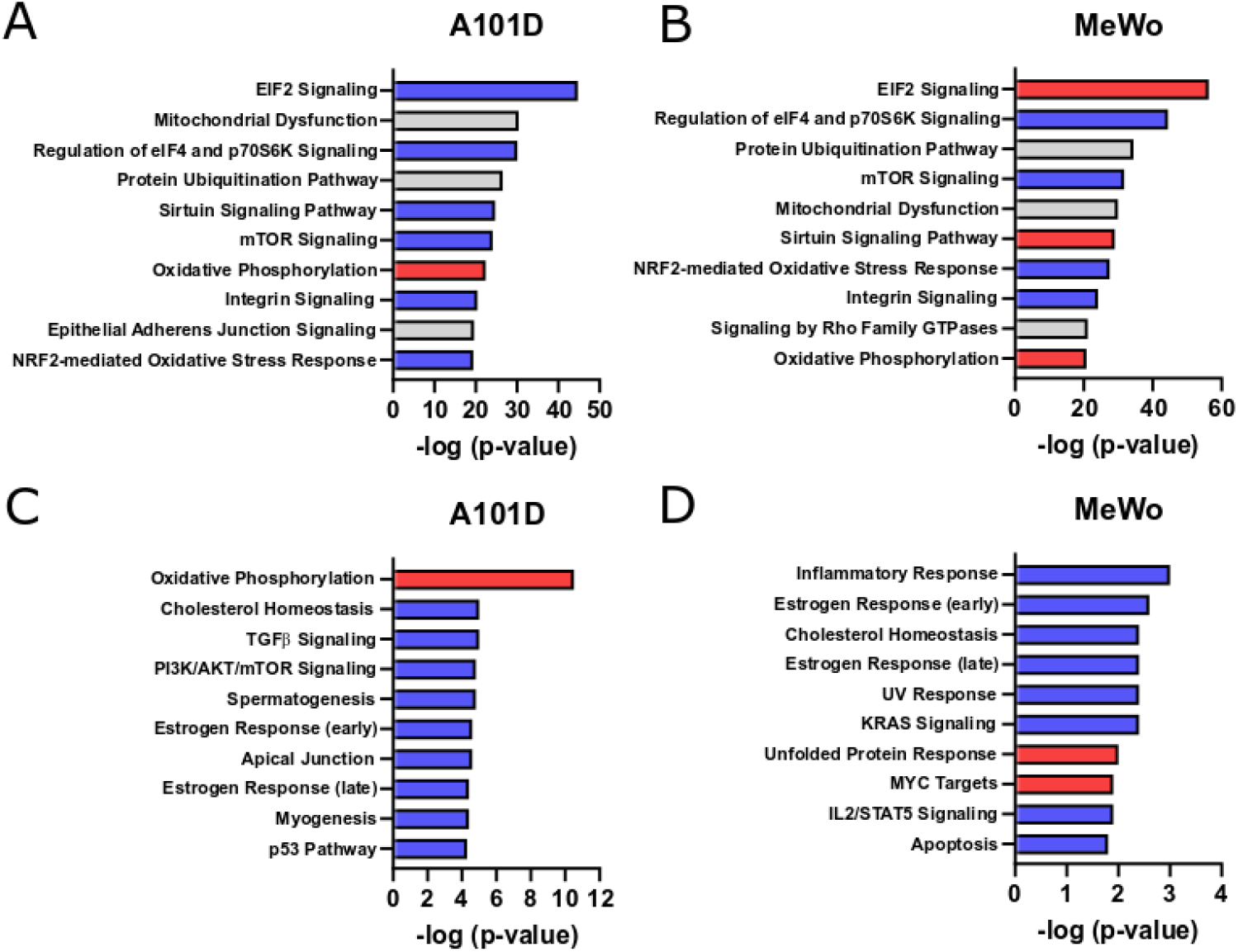
Significant pathways in crude mitochondrial extract. Top 10 significantly enriched pathways after IPA analysis (A and B) and EGSEA hallmark analysis (C and D) of altered proteins in a crude mitochondrial extract in cells cultured for 24h in methionine-free media supplemented with homocysteine compared to control methionine-containing media in A101D (A and C) and MeWo (B and D). Red bars indicate a positive predicted effect, and blue bars indicate a negative predicted effect.

### ATP production does not distinguish between methionine independent and dependent

To gain insight about oxidative phosphorylation in methionine dependence, we measured cellular oxygen consumption and acidification with a Seahorse system. In both A101D and MeWo, there was a decreased in both oxygen production and acidification with homocysteine compared to methionine (Fig 4, A-D). The basal respiration, total amount of produced ATP, ATP from oxidative phosphorylation, and ATP from glycolysis were all decreased to a similar extent in both cell lines in homocysteine compared to methionine (Figure 4, panels E and F). We conclude that it is unlikely that the difference in survival under homocysteine conditions between A101D and MeWo is due solely to deficiencies in oxidative phosphorylation. We therefore shifted our attention to another possible effect of ATF4 upregulation; ferroptosis.

**Figure 4.**
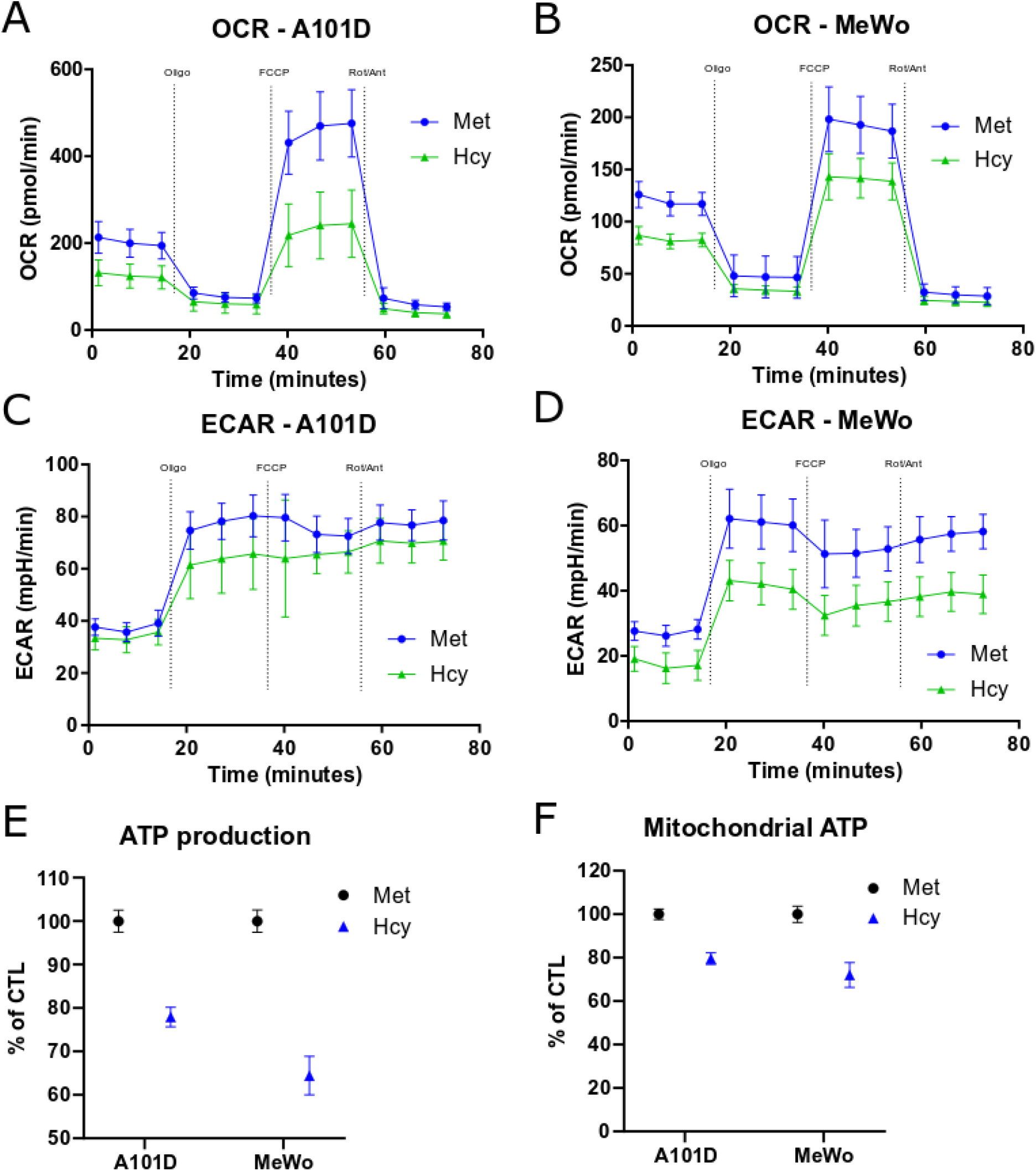
Cellular bioenergetics in cells fed either methionine or homocysteine. Mitochondrial respiration represented by the oxygen consumption rate (OCR) (A and B) and glycolysis represented by the extracellular acidification rate (ECAR) (C and D) in A101D cells (A and C) and MeWo cells (B and D) cultured in media containing methionine or methionine-free media supplemented with homocysteine for 24h. Standard deviation shown. Met: Methionine, Hcy: Methionine-free, homocysteine supplemented medium.

### RNA-Seq analysis reveals a ferroptosis signature associated with homocysteine

To assess the role of ferroptosis in methionine dependence, we analyzed RNA-Seq gene expression data for a ferroptotic cell death signature (30). The median gene expression change for targets in this list was 2.5 in A101D cells grown in homocysteine compared to methionine (Table 2). This value is identical to the one reported by Dixon et al. for the prototypical ferroptosis inducer erastin (30). Median gene change was only 1.27 in MeWo, matching the phenotypic responses observed in the two cell lines. However, the gene and protein expression pattern of iron-related targets were inconsistent with ferroptosis (Table 3). There was a large decrease (4 fold) in the gene expression of both the transferrin receptor and ferroportin in A101D cells grown in homocysteine, while increases have typically been reported with ferroptosis. The transferrin receptor was also decreased at the protein level. Ferritin light and heavy chain, typically degraded through ferritinophagy in ferroptosis, were instead increased at both the gene expression and protein levels (Table 3).

**Table 2.** Ferroptosis gene signature.

RNA-Seq analysis reveals a ferroptosis signature associated with homocysteine
|  | A101D | MeWo |
| --- | --- | --- |
| CHAC1 | 11.60 | 2.60 |
| DDIT4 | 3.53 | 1.33 |
| LOC28456 | nd | nd |
| ASNS | 4.44 | 1.58 |
| TSC22D3 | 2.35 | 1.30 |
| DDIT3 | 3.15 | 1.36 |
| JDP2 | 3.46 | 1.36 |
| SESN2 | 3.48 | 1.59 |
| SLC1A4 | 1.57 | 1.06 |
| PCK2 | 1.74 | 1.33 |
| SLC7A11 | 5.98 | 0.95 |
| TXNIP | 2.27 | 1.14 |
| VLDLR | 1.62 | 1.09 |
| GPT2 | 3.87 | 1.18 |
| PSAT1 | 3.29 | 1.28 |
| C9ORF150 | 4.81 | 1.33 |
| SLC7A5 | 2.04 | 0.98 |
| HERPUD1 | 1.38 | 1.20 |
| XBP1 | 0.95 | 1.06 |
| ATF3 | 2.20 | 1.26 |
| SLC3A2 | 2.25 | 1.01 |
| CBS | 2.43 | 1.21 |
| ATF4 | 2.55 | 1.67 |
| ZNF419 | 2.45 | 1.12 |
| KLHL24 | 1.17 | 1.22 |
| TRIB3 | 5.50 | 1.59 |
| ZNF643 | 1.45 | 0.97 |
| ATP6V1G2 | 2.28 | 1.65 |
| VEGFA | 3.21 | 1.19 |
| GDF15 | 5.79 | 1.38 |
| TUBE1 | 3.05 | 1.43 |
| ARRDC3 | 1.02 | 0.84 |
| CEBPG | 6.66 | 1.51 |
| Median | 2.50 | 1.27 |

**Table 3.** Iron-related targets.

|  | RNA |  | Nuclear |  | Mito |  |
| --- | --- | --- | --- | --- | --- | --- |
|  | MeWo | A101D | MeWo | A101D | MeWo | A101D |
| TFRC | 0.81 | 0.24 | 0.85 | 0.60 | 0.53 | 0.41 |
| SLC40A1 (ferroportin) | 1.73 | 0.24 | undet | undet | undet | undet |
| FTH1 | 0.95 | 1.39 | 1.58 | 2.28 | 1.57 | 1.46 |
| FTL | 0.97 | 1.43 | 1.55 | 2.69 | 1.71 | 1.16 |

### Homocysteine induces lipid peroxidation in methionine dependent cells

To confirm the role of ferroptotic cell death in methionine dependence, we measured the fluorescence shift of BODIPY dye from 581 to 591nm as a marker of lipid peroxidation. After 24h, there was a significant increase in the level of emission at 591nm over 581nm detected in A101D cells grown in homocysteine media (Fig 5, A and B). This increase was not due to the presence of homocysteine itself, and was not detected in the methionine independent cell line MeWo.

**Figure 5.**
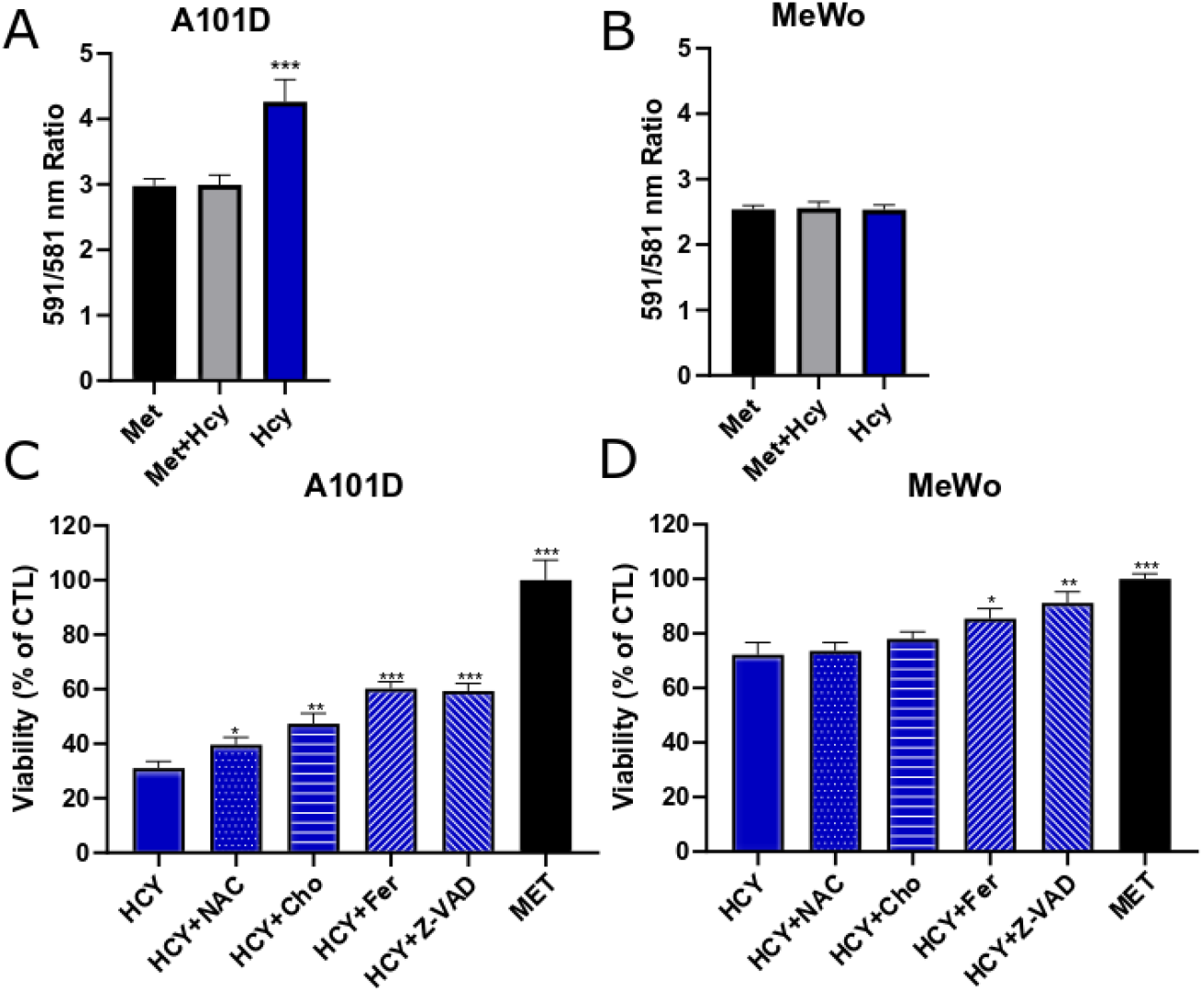
Lipid peroxidation in cells fed either methionine or homocysteine. BODIPY intensity levels in A101D (A) and MeWo (B) cells cultured in media containing methionine, methionine and homocysteine, or methionine-free media supplemented with homocysteine for 24h. Cell viability/proliferation assessed with resazurin in A101D (C) and MeWo (D) cells cultured in media containing methionine or methionine-free media supplemented with homocysteine for 72h. MET: Methionine-containing media, HCY: Methionine-free, homocysteine supplemented medium, NAC: N-acetylcysteine, Cho: choline, Fer: Ferrostatin, Z-VAD: Z-VAD-FMK.

### Ferrostatin rescues methionine dependent cells

Finally, we sought to clarify further the cause of death in methionine dependent cells by attempting to rescue cells with a panel of compounds. Both ferrostatin and Z-VAD-FMK rescued viability in methionine dependent A101D cells grown in homocysteine (Fig 5, C and D). Ferrostatin also rescued viability in the murine melanoma cell line B16F10 (Supplementary Fig 2). Methionine can provide methyl groups for the synthesis of phosphatidylcholine for cellular membranes. However, viability was only modestly rescued by supplementation with choline.

## DISCUSSION

Cancer cells are generally more resistant to harsh environmental conditions than normal cells. However, they also display specific vulnerabilities. Methionine dependence in cancer cells is one of such vulnerabilities, described over 45 years ago. We attempted to characterize the molecular bases behind the inability of most cancer cells to utilize homocysteine instead of methionine to sustain growth and metabolism.

In order to validate our system, we performed an analysis of the cell cycle in a methionine independent and a methionine dependent cell line. As expected, proliferation was decreased at both 24 and 48h in A101D cells, in contrast to a slight increase cells undergoing mitosis in MeWo. An increase in cells in subG1 phase in A101D cells after 48h suggested an apoptotic mechanism of cell death in methionine dependent cells grown in methionine-free, homocysteine-containing media.

A single methyl group differentiates methionine from homocysteine. We therefore looked at differences in DNA methylation between A101D and MeWo. DNA hypomethylation was a hallmark in both cell lines, but interestingly, the effect was more profound in MeWo cells. Continued proliferation in these cells may lead to a dilution effect compared to non-proliferating A101D cells. Although some signaling pathways were significantly enriched, both positive and negative regulators were hypomethylated, leading to an uncertain outcome on cellular function. An insight into function came from the analysis of the nuclear proteome, which indicated an activation of the mitochondrial stress response (31). Although detectable in both cell lines, it was more pronounced in methionine dependent cells compared to methionine independent cells. The increase in the nuclear abundance of the master regulator of the mitochondrial stress response ATF4 observed in A101D was also consistent with the activation of pathways to increase the biosynthesis of amino acids (31). An increase in ATF4 gene expression has previously been identified in methionine-restricted triple-negative breast cancer cells (32). It was reported to induce the integrated stress response independently of the kinases GCN2 and PERK (32).

The analysis of a fraction enriched for mitochondrial proteins indicated changes in oxidative phosphorylation. The oxygen consumption rate of A101D cells grown in homocysteine versus methionine was indeed lower. However, this decrease was comparable to the one measured in MeWo, in which proliferation is unaffected, except for the maximal respiratory rate. ATF4 is however also a marker of ferroptotic cell death. We identified a strong gene signature for ferroptosis in A101D, which was present at a lower level in MeWo. Consistent with ferroptotic cell death, there was a small but significant increase in lipid peroxidation in A101D, and viability could be partially rescued with the inhibitor of ferroptosis ferrostatin. Interestingly, the caspase inhibitor Z-VAD-FMK also rescued cell viability. This confirms the role of ferroptosis in death in methionine dependence. In this model, ferroptotic cell death was happening in conjunction with apoptotic cell death, as suggested initially by the increase in subG1 cells. The two mechanisms, once believed to be mutually exclusive, have been reported to occur concomitantly (33). The balance between apoptosis and ferroptosis is likely to be influenced by cell density, as previously suggested (34). Future experiments will assess whether these two forms of cell death are also relevant to *in vivo* tumors.

## ACKNOWLEDGEMENTS

The authors would like to thank Giulia Baldini and Nükhet Aykin-Burns for guidance, as well as Robin Mulkey and other employees at the UAMS animal facility for their assistance. Kathy Bush provided emergency childcare during the COVID crisis. The UAMS Bioinformatics core is supported by the Winthrop P. Rockefeller Cancer Institute and National Institutes of Health grant P20GM121293. This work was supported by the Winthrop P. Rockefeller Cancer Institute, the Arkansas Tobacco Settlement Commission, and by the National Center for Advancing Translational Sciences grant number 1KL2TR003108-01.

## SUPPLEMENTARY FIGURES

**Supplementary Figure 1.**
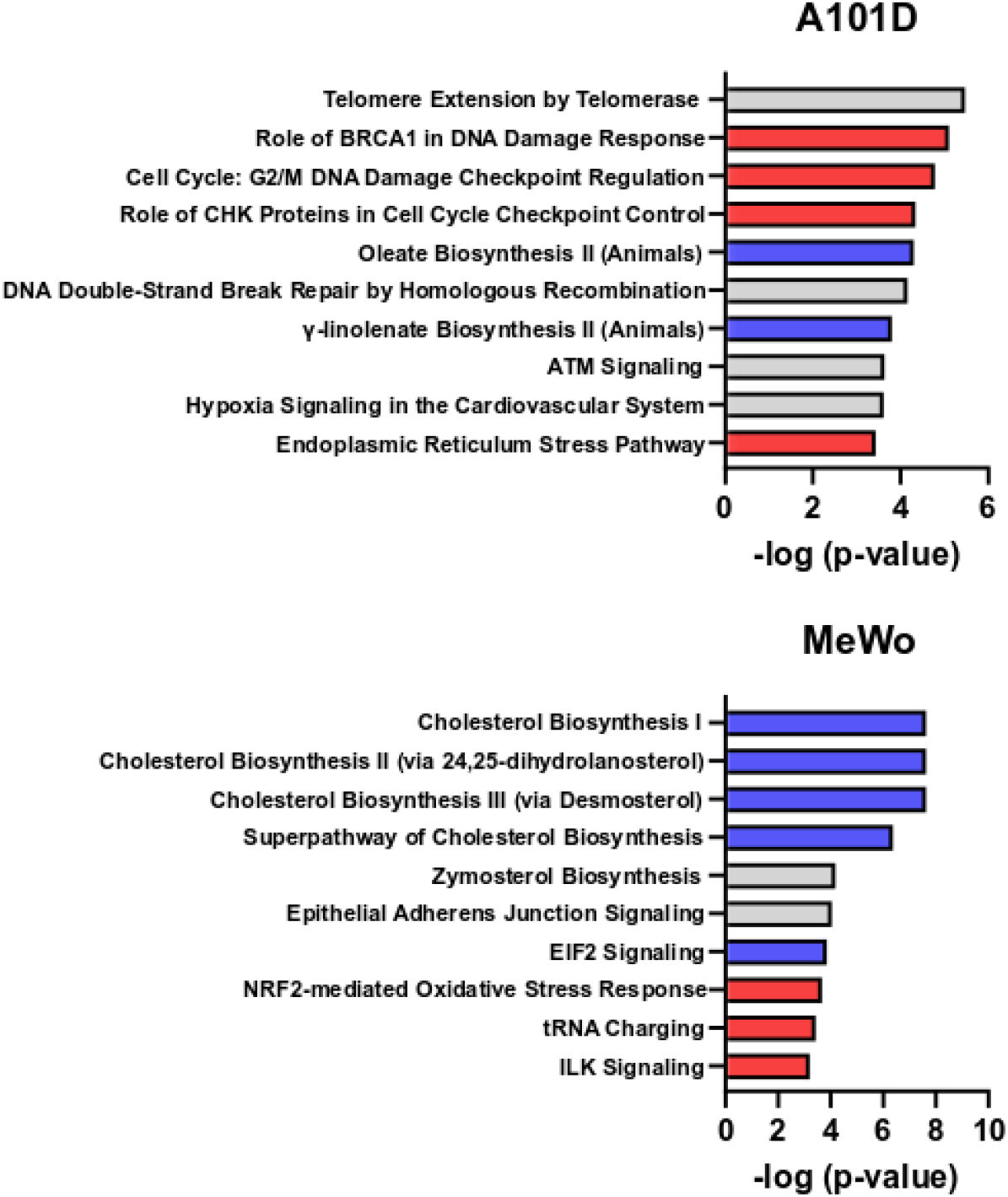
Significant pathways in nuclear extract. Top 10 pathways enriched after IPA analysis of nuclear proteins in A101D and MeWo cells cultured in media containing methionine or methionine-free media supplemented with homocysteine for 24h. Red bars indicate a positive predicted effect, and blue bars indicate a negative predicted effect.

**Supplementary Figure 2.**
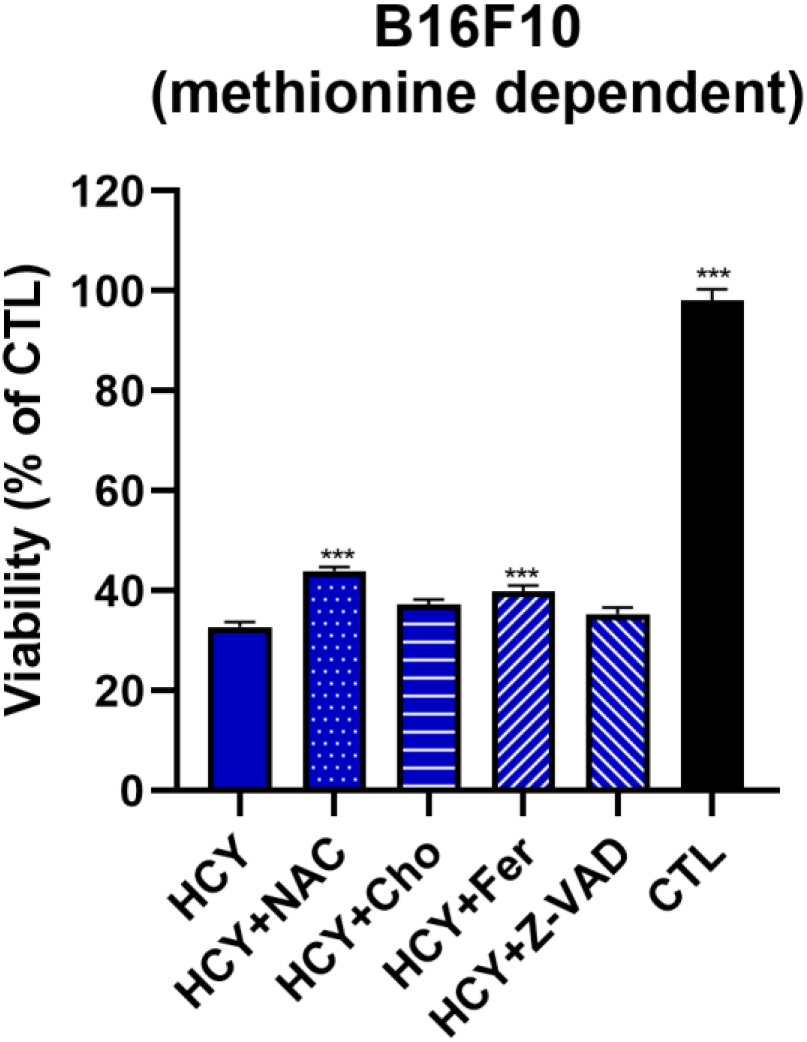
Cell viability in B16F10 murine melanoma cells. Cell viability/proliferation assessed with resazurin in B16F10 murine melanoma cells cultured in media containing methionine or methionine-free media supplemented with homocysteine for 72h. CTL: Methionine-containing media, HCY: Methionine-free, homocysteine supplemented medium, NAC: N-acetylcysteine, Cho: choline, Fer: Ferrostatin, Z-VAD: Z-VAD-FMK.

